# Assembly and Annotation of an Ashkenazi Human Reference Genome

**DOI:** 10.1101/2020.03.18.997395

**Authors:** Alaina Shumate, Aleksey V. Zimin, Rachel M. Sherman, Daniela Puiu, Justin M. Wagner, Nathan D. Olson, Mihaela Pertea, Marc L. Salit, Justin M. Zook, Steven L. Salzberg

**Affiliations:** Center for Computational Biology, Johns Hopkins University, Baltimore, MD; Department of Biomedical Engineering, Johns Hopkins University, Baltimore, MD; Department of Computer Science, Johns Hopkins University, Baltimore, MD; National Institute of Standards and Technology, Gaithersburg, MD; Joint Initiative for Metrology in Biology, Stanford University, Stanford, CA; Department of Biostatistics, Johns Hopkins University, Baltimore, MD

**Author notes:** These authors contributed equally to this work (Alaina Shumate, Aleksey V. Zimin).

## Abstract

Here we describe the assembly and annotation of the genome of an Ashkenazi individual and the creation of a new, population-specific human reference genome. This genome is more contiguous and more complete than GRCh38, the latest version of the human reference genome, and is annotated with highly similar gene content. The Ashkenazi reference genome, Ash1, contains 2,973,118,650 nucleotides as compared to 2,937,639,212 in GRCh38. Annotation identified 20,157 protein-coding genes, of which 19,563 are >99% identical to their counterparts on GRCh38. Most of the remaining genes have small differences. 40 of the protein-coding genes in GRCh38 are missing from Ash1; however, all of these genes are members of multi-gene families for which Ash1 contains other copies. 11 genes appear on different chromosomes from their homologs in GRCh38. Alignment of DNA sequences from an unrelated Ashkenazi individual to Ash1 identified ~1 million fewer homozygous SNPs than alignment of those same sequences to the more-distant GRCh38 genome, illustrating one of the benefits of population-specific reference genomes.

## Introduction

The human reference genome is used as a resource for many thousands of experiments and studies every year. Since 2001, the international community has relied on a single reference genome (currently GRCh38) that is a mosaic of sequence from a small number of individuals, with about 65% originating from a single person (Green et al. 2010), who was later identified as being approximately 50% European and 50% African by descent. The current 3-gigabase reference sequence is a vastly improved version of the genome that was published in 2001 (International Human Genome Sequencing Consortium 2001), but it represents a miniscule sample of the human population, currently estimated at just under 8 billion people. In the future, the scientific community will likely have hundreds and eventually thousands of reference genomes, representing many different sub-populations. For now, though, all human protein-coding genes, RNA genes, and other important genetic features are mapped onto the coordinate system of the reference genome, as are millions of single-nucleotide polymorphisms (SNPs) and larger structural variants. Large-scale SNP genotyping arrays, exome capture kits, and countless other genetic analysis tools are also based on GRCh38.

Many studies have pointed out that a single genome is inadequate for a variety of reasons, such as inherent bias towards the reference genome (Need and Goldstein 2009, Popejoy and Fullerton 2016, Ballouz et al. 2019). The availability of reference genomes from multiple human populations would greatly aid attempts to find genetic causes of traits that are over- or under-represented in those populations, including susceptibility to disease (Wong et al. 2018). Another drawback of relying on a single reference genome is that it almost certainly contains minor alleles at some loci, which in turn confounds studies focused on variant discovery and association of those variants with disease (Ferrarini et al. 2015, Magi et al. 2015, Barbitoff et al. 2018, Wong et al. 2018).

The worldwide scientific community is currently engaged in whole-genome sequencing of hundreds of thousands of people, and several countries have announced plans to sequence millions more. Despite this enormous investment, the initial analysis of all of these genomes relies, for now, on just one reference genome, GRCh38. Variants present in regions that are missing from this genome will be essentially invisible until other reference genomes are available. Although many human genome assemblies have been published in recent years, none has undergone the full set of steps, particularly annotation, necessary to create a reference genome that can be used in the same manner as GRCh38 (although the Korean AK1 genome (Seo et al. 2016) included some annotation). Necessary steps include ordering and orienting all contigs along chromosomes, filling in gaps as much as possible, and annotating the resulting assembly with all known human genes. Because so much of the literature also relies on the current naming system for human genes, annotation of new reference genomes should also use the same terminology and gene identifiers to be maximally useful. Here we describe the first such effort to create an alternative human reference genome, which we have called Ash1, based on deep sequencing of an Ashkenazi individual. The Ash1 genome and annotation is freely available through https://github.com/AshkenaziGenome/Assembly, and has been deposited in GenBank as accession GCA_011064465.1 and BioProject PRJNA607914.

## Results

For creation of the first human reference genome to be assembled from a single individual, we chose HG002, an Ashkenazi individual who is part of the Personal Genome Project (PGP). The PGP uses the Open Consent Model, the first truly openaccess platform for sharing individual human genome, phenotype, and medical data (Church 2005, Ball et al. 2014). The consent process educates potential participants on the implications and risks of sharing genomic data, and about what they can expect from their participation. Open consent has allowed for the creation of the world’s first human genome reference materials (HG002 is NIST Reference Material 8391) from Genome In A Bottle (GIAB), which is being used for calibration, genome assembly methods development, and lab performance measurements (Zook et al. 2014, Zook et al. 2019). All raw sequence data for this project was obtained from GIAB, where it is freely available to the public (Zook et al. 2016).

We assembled the HG002 genome from a combination of three deep-coverage data sets: 249-bp Illumina reads, Oxford Nanopore (ONT) reads averaging over 33 Kbp in length, and high-quality PacBio “HiFi” reads averaging 9567 bp (Table 1).

**Table 1.** Sequence data for assembly of the HG002 genome, all taken from the Genome In A Bottle Project.

|  |  |  |  |  |
| --- | --- | --- | --- | --- |
| **Sequencing Technology** | **Number of reads** | **Mean read length (bp)** | **Total sequence (bp)** | **Genome coverage** |
| **Illumina** | 883,914,482 | 249 | 219,763,641,914 | 71x |
| **ONT** | 2,090,962 | 33,889 | 70,861,178,054 | 23x |
| **PacBio HiFi** | 9,270,502 | 9,567 | 88,695,245,383 | 29x |

We initially created two assemblies, one using Illumina and ONT reads, and a second using all three data sets, including the PacBio HiFi reads. The addition of PacBio HiFi data resulted in slightly more total sequence in the assembly (2.99 Gb vs. 2.88 Gb) and a substantially larger contig N50 size (18.2 Mb vs 4.9 Mb). This assembly, designated Ash1 v0.5, was the basis for all subsequent refinements.

### Mapping the assembly onto chromosomes

To create chromosome assignments for the Ash1 v0.5 assembly, we used alignments to GRCh38 to map most of the scaffolds onto chromosomes. The steps described in **Methods** generated a series of gradually improved chromosome-scale assemblies, resulting in Ash1 v1.7. Ash1 v1.7 has greater contiguity and smaller gaps than GRCh38, as shown in Table 2. Note that in the process of building these chromosomes, a small amount of GRCh38 sequence (58.3 Mb, 2% of the genome) was used to fill gaps in Ash1. These regions include some difficult-to-assemble regions that have been manually curated for GRCh38. In total, the estimated size of all gaps in Ash1 is 82.9 Mbp, versus 84.7 Mbp in GRCh38.p13.

**Table 2.**
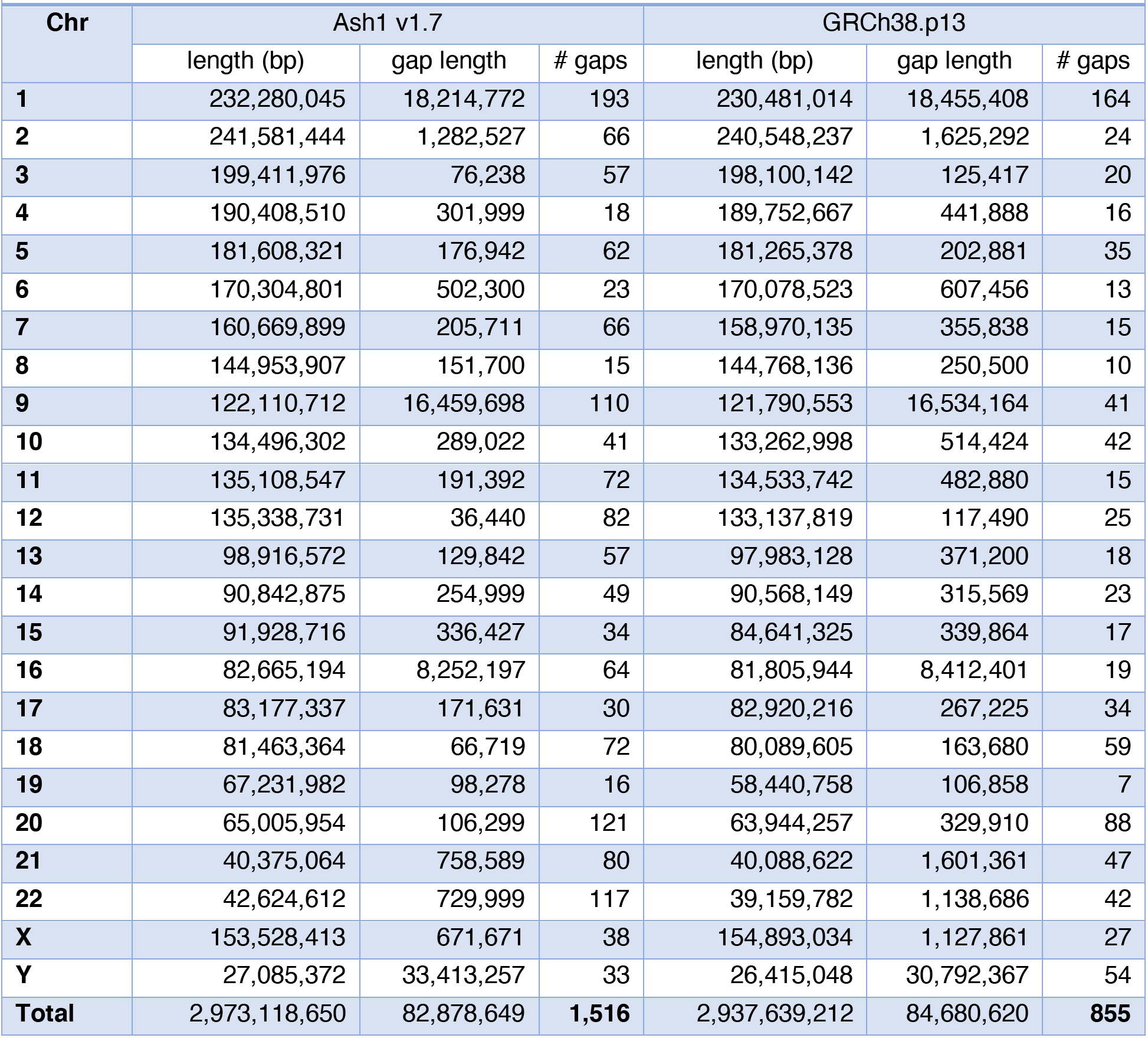
Comparison of chromosome lengths and gaps between Ash1 and GRCh38. Chromosome lengths exclude all “N” characters. Every sequence of Ns was counted as a gap except for leading and trailing Ns. Several GRCh38 chromosomes begin or end with lengthy sequences of Ns numbering millions of bases; these were not counted as gaps here.

| Chr | Ash1 v1.7 |  |  | GRCh38.p13 |  |  |
| --- | --- | --- | --- | --- | --- | --- |
|  | length (bp) | gap length | # gaps | length (bp) | gap length | # gaps |
| 1 | 232,280,045 | 18,214,772 | 193 | 230,481,014 | 18,455,408 | 164 |
| 2 | 241,581,444 | 1,282,527 | 66 | 240,548,237 | 1,625,292 | 24 |
| 3 | 199,411,976 | 76,238 | 57 | 198,100,142 | 125,417 | 20 |
| 4 | 190,408,510 | 301,999 | 18 | 189,752,667 | 441,888 | 16 |
| 5 | 181,608,321 | 176,942 | 62 | 181,265,378 | 202,881 | 35 |
| 6 | 170,304,801 | 502,300 | 23 | 170,078,523 | 607,456 | 13 |
| 7 | 160,669,899 | 205,711 | 66 | 158,970,135 | 355,838 | 15 |
| 8 | 144,953,907 | 151,700 | 15 | 144,768,136 | 250,500 | 10 |
| 9 | 122,110,712 | 16,459,698 | 110 | 121,790,553 | 16,534,164 | 41 |
| 10 | 134,496,302 | 289,022 | 41 | 133,262,998 | 514,424 | 42 |
| 11 | 135,108,547 | 191,392 | 72 | 134,533,742 | 482,880 | 15 |
| 12 | 135,338,731 | 36,440 | 82 | 133,137,819 | 117,490 | 25 |
| 13 | 98,916,572 | 129,842 | 57 | 97,983,128 | 371,200 | 18 |
| 14 | 90,842,875 | 254,999 | 49 | 90,568,149 | 315,569 | 23 |
| 15 | 91,928,716 | 336,427 | 34 | 84,641,325 | 339,864 | 17 |
| 16 | 82,665,194 | 8,252,197 | 64 | 81,805,944 | 8,412,401 | 19 |
| 17 | 83,177,337 | 171,631 | 30 | 82,920,216 | 267,225 | 34 |
| 18 | 81,463,364 | 66,719 | 72 | 80,089,605 | 163,680 | 59 |
| 19 | 67,231,982 | 98,278 | 16 | 58,440,758 | 106,858 | 7 |
| 20 | 65,005,954 | 106,299 | 121 | 63,944,257 | 329,910 | 88 |
| 21 | 40,375,064 | 758,589 | 80 | 40,088,622 | 1,601,361 | 47 |
| 22 | 42,624,612 | 729,999 | 117 | 39,159,782 | 1,138,686 | 42 |
| X | 153,528,413 | 671,671 | 38 | 154,893,034 | 1,127,861 | 27 |
| Y | 27,085,372 | 33,413,257 | 33 | 26,415,048 | 30,792,367 | 54 |
| **Total** | **2,973,118,650** | **82,878,649** | **1,516** | **2,937,639,212** | **84,680,620** | **855** |

As part of the assembly improvement process, we searched one of the preliminary Ash1 assemblies (v1.1) for the 12,745 high-quality, isolated structural variants (insertions and deletions ≥50 bp) that Zook et al. identified by comparing the Ashkenazi trio data to GRCh37 (Zook et al. 2019). That study used four different sequencing technologies and multiple variant callers to identify variants and filter out false positives. Of these 12,745 SVs, 5807 are homozygous and 6938 are heterozygous. We expected the Ash1 assembly to agree with nearly all of the homozygous variants. Because Ash1 captures just one haplotype, we expected that it would agree with approximately half of the heterozygous SVs, assuming that the assembly algorithm chose randomly between the haplotypes when deciding which variant to include in the final consensus. Of the 5807 homozygous variants, 5284 (91%) were present using our match criteria (see Methods), and 3922 (56.5%) of 6938 heterozygous variants were present. All variants were found at the correct location.

We also made small (≤5bp) variant calls on Ash1 v1.1 and compared these to the HG002 v4.0 benchmark variants from GIAB, which we used to correct numerous substitution and indel errors (see Methods), yielding Ash 1 v1.2. We then re-aligned the Ash1 assembly to GRCh38, re-called variants, and benchmarked these variants against the newly-developed v4.1 GIAB benchmark set. Of the variants inside the v4.1 benchmark regions, the Ash1 variants matched 1,256,458 homozygous and 1,041,476 heterozygous SNPs, and 187,227 homozygous and 193,524 heterozygous indels. After excluding variant calls within 30bp of a true variant, 79,269 SNPs and 17,439 indels remained, which (assuming these are all errors in Ash1) corresponds to a quality value (QV) of approximately Q45 for substitution errors. Most of these variants (52,191 SNPs and 4629 indels) fall in segmental duplications, possibly representing missing duplications in Ash1 or imperfect polishing by short reads. In summary, the quality of the Ash1 assembly is very high, with an estimated substitution quality value of 62 and an indel error rate of 2 per million bp after excluding known segmental duplications, tandem repeats, and homopolymers.

### Comparison of variant calling using Ash1 versus GRCh38

One of the motivations for creating new reference genomes is that they provide a better framework for analyzing human sequence data when searching for genetic variants linked to disease. For example, a study of Ashkenazi Jews that collected whole-genome shotgun (WGS) data should use an Ashkenazi reference genome rather than GRCh38. Because the genetic background is similar, fewer variants should be found when searching against Ash1, and the variants that do appear will be more likely to be disease-relevant.

To test this expectation, we collected WGS data from a male participant in the Personal Genome Project, PGP17 (hu34D5B9). This individual is estimated to be 66% Ashkenazi according to the PGP database, which was the highest estimated fraction available from already-sequenced PGP individuals. We downloaded 983,220,918 100-bp reads (approximately 33x coverage) and aligned them to both Ash1 and GRCh38 using Bowtie2 (Langmead and Salzberg 2012). Slightly more reads (3,901,270, 0.5%) aligned to Ash1 than to GRCh38.

We then examined all single-nucleotide variants (SNVs, see Methods) between PGP17 and each of the two reference genomes. To simplify the analysis, we only considered locations where PGP17 was homozygous. In our comparisons to Ash1, we first identified all SNVs, and then examined the original Ash1 read data to determine whether, for each of those SNVs, the Ash1 genome contained a different allele that matched PGP17.

In total, the number of homozygous sites in PGP17 that disagreed with Ash1 was 1,333,345, versus 1,700,364 when we compared homozygous sites in PGP17 to GRCh38 (**Supplementary Table S1**). We then looked at the underlying Ash1 read data for the 1.33 million SNV sites that initially mismatched, and found that for approximately half of them, the Ash1 genome was heterozygous, with one allele matching PGP17. If we restricted SNVs to sites where PGP17 and Ash1 are both homozygous (plus a very small number of locations where Ash1 contains two variants that both differ from PGP17) this reduced the total number of SNVs even further, to 707,756. Thus we found just under 1 million fewer homozygous SNVs when using Ash1 as the reference for PGP17.

### Comparison against common Ashkenazi variants

To examine the extent to which Ash1 contains known, common Ashkenazi variants (relative to GRCh38), we examined SNVs reported at high frequency in an Ashkenazi population from the Genome Aggregation Database (gnomAD) (Karczewski et al. 2019). GnomAD v3.0 contains SNV calls from short-read whole-genome data from 1,662 Ashkenazi individuals. Because some variants were only called in a subset of these individuals, we considered only variant sites that were reported in a minimum of 200 people. We then collected major allele SNVs, requiring the allele frequency to be above 0.5 in the sampled population. We further restricted our analysis to single-base substitutions. This gave us 2,008,397 gnomAD SNV sites where the Ashkenazi major allele differed from GRCh38.

We were able to precisely map 1,790,688 of the 2,008,397 gnomAD sites from GRCh38 onto Ash1 (see Methods). We then compared the GRCh38 base to the Ashkenazi major allele base at each position, and we also examined the alternative alleles in Ash1 at sites where it is heterozygous. For sites where the read data showed that HG002 was heterozygous and had both the Ashkenazi major allele and the GRCh38 allele, we replaced the Ash1 base, if necessary, to ensure that it matched the major allele. After these replacements, Ash1 contained the Ashkenazi major allele at 88% (1,580,866) of the 1.79 million sites. At the remaining sites, Ash1 either matched the GRCh38 allele because HG002 is homozygous for the reference allele (204,729 sites), or it contained a third allele matching neither GRCh38 nor the gnomAD major allele (5,093 sites).

Worth noting is that, as the frequency of the major allele in the gnomAD Ashkenazi population increases, the proportion of sites where Ash1 matched the major allele increases as well. For example, of SNVs that have an allele frequency >0.9 in the gnomAD Ashkenazi population, over 98% are represented in Ash1 (**Table 3**).

**Table 3.** The proportion of variant sites in the Ashkenazi reference genome that agree with major alleles from the gnomAD large-scale survey of the Ashkenazi population. Column headers show the frequency ranges of the Ashkenazi alternative alleles (ALT) from the gnomAD database. Row 3 shows the proportion of positions in Ash1 that agree with the gnomAD major allele where gnomAD differs from GRCh38.

| Frequency (f) in Ashkenazi population | [0.25, 0.5] | (0.5, 0.6] | (0.6, 0.7] | (0.7, 0.8] | (0.8, 0.9] | (0.9, 1.0] | Total |
| --- | --- | --- | --- | --- | --- | --- | --- |
| Total # sites at Ashkenazi ALT allele frequency (f) | 1,706,379 | 442,352 | 369,541 | 300,969 | 252,859 | 424,967 | **3,497,067** |
| Proportion of Ash1 sites that match gnomAD Ashkenazi allele | 0.317 | 0.759 | 0.846 | 0.910 | 0.955 | 0.982 | **0.607** |

### Annotation

A vital part of any reference genome is annotation: the collection of all genes and other features found on the genome. To allow Ash1 to function as a true reference genome, we have mapped all of the known genes used by the scientific community onto its coordinate system, using the same gene names and identifiers. Several different annotation databases have been created for GRCh38, and for Ash1 we elected to use the CHESS human gene database (Pertea et al. 2018) because it is comprehensive, including all of the protein-coding genes in both GENCODE and RefSeq, the two other widely-used gene databases, and because it retains the identifiers used in those catalogs. The noncoding genes differ among the three databases, but CHESS has the largest number of gene loci and isoforms. We used CHESS version 2.2, which has 42,167 genes on the primary chromosomes (excluding the GRCh38 alternative scaffolds), of which 20,197 are protein coding.

Mapping genes from one assembly to another is a complex task, particularly for genes that occur in highly similar, multi-copy gene families. The task is even more complex when the two assemblies represent different individuals (rather than simply different assemblies of the same individual), due to the presence of single-nucleotide differences, insertions, deletions, rearrangements, and genuine variations in copy number between the individuals. We needed a method that would be robust in the face of all of these potential differences.

To address this problem, we used the recently developed Liftoff mapping tool, which in our experiments was the only tool that could map nearly all human genes from one individual to another. Liftoff takes all of the genes and transcripts from a genome and maps them, chromosome by chromosome, to a different genome. For all genes that fail to map to the same chromosome, Liftoff attempts to map them across chromosomes. Unlike other tools, it does not rely on a detailed map built from a whole-genome alignment, but instead it maps each gene individually. Genes are aligned at the transcript level, including introns, so that processed pseudogenes will not be mistakenly identified as genes.

We attempted to map all 310,901 transcripts from 42,167 gene loci on the primary chromosomes in GRCh38 to Ash1. In total, we successfully mapped 309,900 (99.7%) transcripts from 42,070 gene loci onto the main chromosomes (**Supplementary Table 2**). We considered a transcript to be mapped successfully if the mRNA sequence in Ash1 is at least 50% as long as the mRNA sequence on GRCh38. However, the vast majority of transcripts greatly exceed this threshold, with 99% of transcripts mapping at a coverage greater than or equal to 95% (**Suppl. Fig S2**). The sequence identity of the mapped transcripts is similarly high, with 99% of transcripts mapping with a sequence identity greater than or equal to 94% (**Suppl. Fig S3**).

#### Translocated genes

Of those genes with at least one successfully mapped isoform, 42,059 (99.7%) mapped to the corresponding locations on the same chromosome in Ash1. Of the 108 genes that initially failed to map, 11 genes mapped to a different chromosome in 7 distinct blocks (shown in **Table 4**), suggesting a translocation between the two genomes. Interestingly, 16 of the 22 locations involved in the translocations were in sub-telomeric regions, which occurred in 8 pairs where both locations were near telomeres. This is consistent with previous studies reporting that rearrangements involving telomeres and subtelomeres may be a common form of translocation in humans (Bailey and Murnane 2006, Liddiard et al. 2016, Muraki and Murnane 2018).

**Table 4.** 11 genes from GRCh38, 4 of them protein coding, that map to a different chromosome on Ash1. Genes are sorted by their position on GRCh38. Genes that appear to have moved in a block via a single translocation are highlighted in colored rows. Sub-telomeric coordinates are indicated by **(T)** next to the coordinates. Abbreviations: NC, noncoding.

| CHESS ID | Gene Name | Gene Type | GRCh38 Location | Ash1 Location |
| --- | --- | --- | --- | --- |
| **CHS.460** | HNRNPCL4 | protein | chr1:13164555-13165482 | chr6:113726526-113727453 |
| **CHS.39870** | USP17L11 | protein | chr4:9215405-9216997 | chr11:71983132-71984724 |
| **CHS.39871** | USP17L12 | protein | chr4:9220152-9221744 | chr11:71978387-71979979 |
| **CHS.54932** | WASH1 | protein | chr9:14475-30487 (T) | chr20:50732-69104 (T) |
| **CHS.54933** | LOC107987041 | NC | chr9:27657-30891 (T) | chr20:65950-69493 (T) |
| **CHS.54934** | FAM138C | NC | chr9:34394-35864 (T) | chr20:65083816-65085286 (T) |
| **CHS.18492** | Unnamed | NC | chr15:101959848-101960582 (T) | chr20:65088782-65089512 (T) |
| **CHS.18493** | WASH3P | NC | chr15:101960813-101976605 (T) | chr20:65089741-65105526 (T) |
| **CHS.18494** | DDX11L9 | NC | chr15:101976558-101979093 (T) | chr20:65105479-65108014 (T) |
| **CHS.20775** | LOC107987240 | NC | chr16:90199813-90211886 (T) | chr20:2-12021 (T) |
| **CHS.59387** | DDX11L16 | NC | chrY:57212178-57214703 (T) | chr20:48248-50782 (T) |

We examined the translocation between chromosomes 15 and 20, which contains three of the genes in Table 4, by looking more closely at the alignment between GRCh38 and Ash1. The translocation is at the telomere of both chromosomes, from position 65,079,275–65,109,824 (30,549 bp) of Ash1 chr20 and 101,950,338–101,980,928 (30,590 bp) of GRCh39 chr15. To confirm the translocation, we aligned an independent set of very long PacBio reads, all from HG002, to the Ash1 v1.7 assembly (see Methods) and evaluated the region around the breakpoint on chr20. These alignments show deep, consistent coverage extending many kilobases on both sides of the breakpoint, supporting the correctness of the Ash1 assembly (Figure 1).

**Figure 1.**
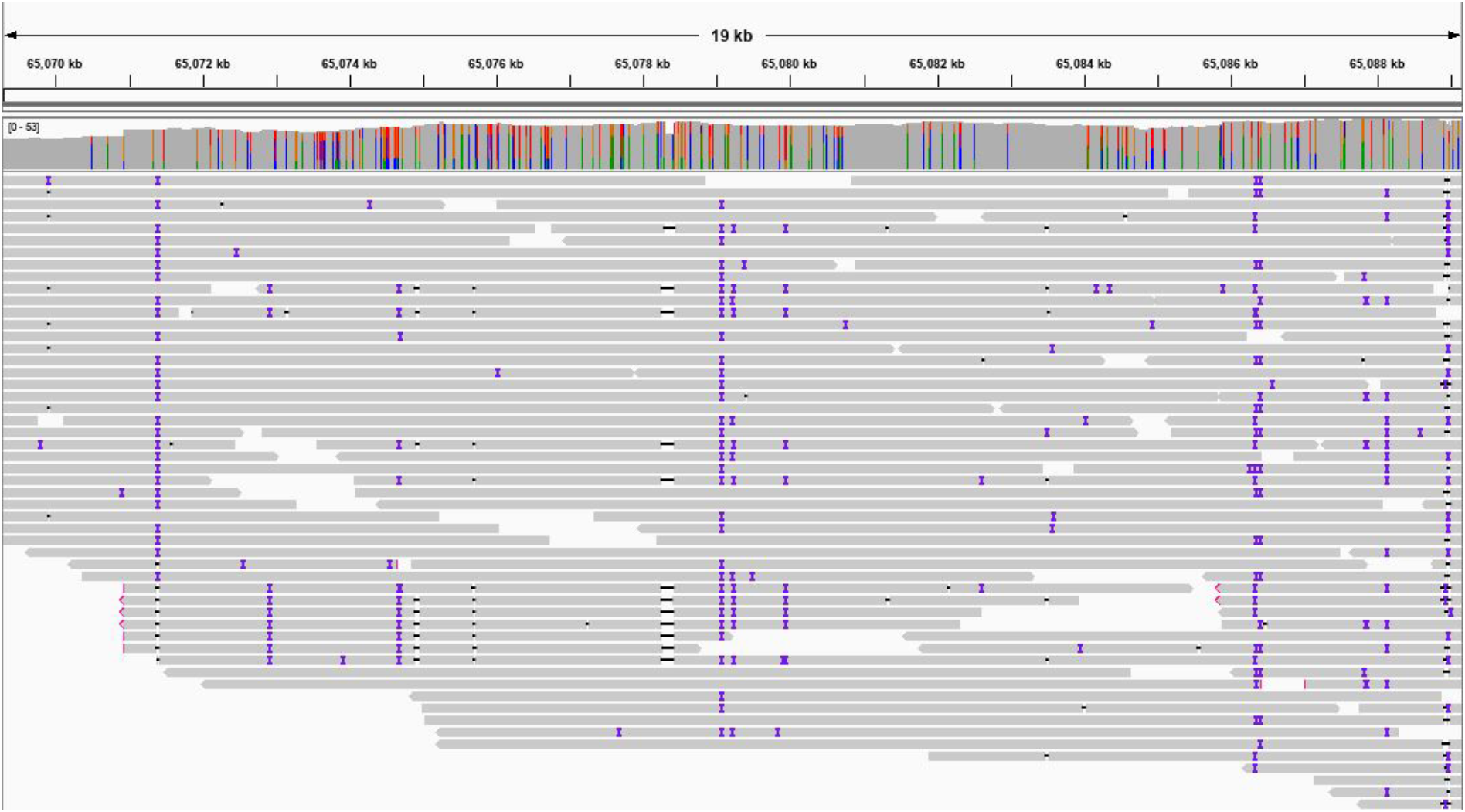
Snapshot showing alignments of long PacBio reads to the Ash1 genome, centered on the left end of the location in chromosome 20 (position 65,079,275) where a translocation occurred between chromosome 15 (GRCh38) and 20 (Ash1). The top portion of the figure shows the coordinates on chr20. Below that is a histogram of read coverage, and the individual reads fill the bottom part of the figure. The indels in the reads, shown as colored bars on each read, are due to the relatively high error rate of the long reads.

#### Missing genes

62 genes failed entirely to map from GRCh38 onto Ash1, and another 32 genes mapped only partially (below the 50% coverage threshold), as shown in **Table 5**. All of the genes that failed to map or that mapped partially were members of multigene families, and in every case there was at least one other copy of the missing gene present in Ash1, at an average identity of 98.5%. Thus there are no cases at all of a gene that is present in GRCh38 and that is entirely absent from Ash1; the genes shown in **Table 5** represent cases where Ash1 has fewer members of a multi-gene family. Three additional genes (2 protein coding, 1 lncRNA) mapped to two unplaced contigs, which will provide a guide to placing those contigs in future releases of the Ash1 assembly.

**Table 5.**
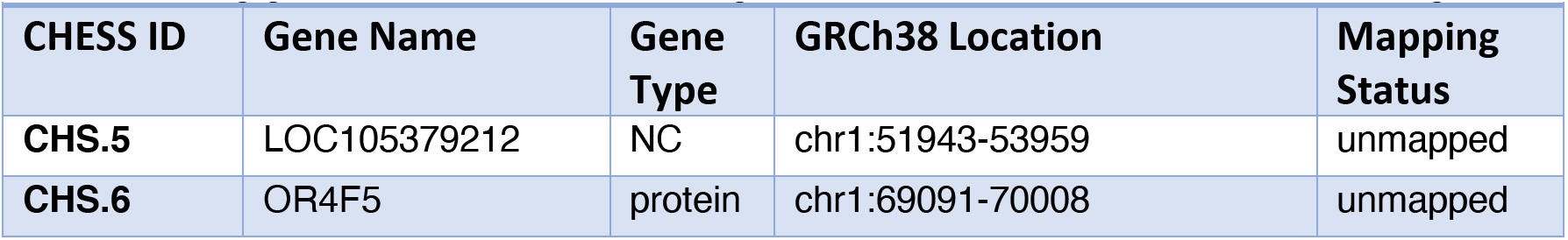
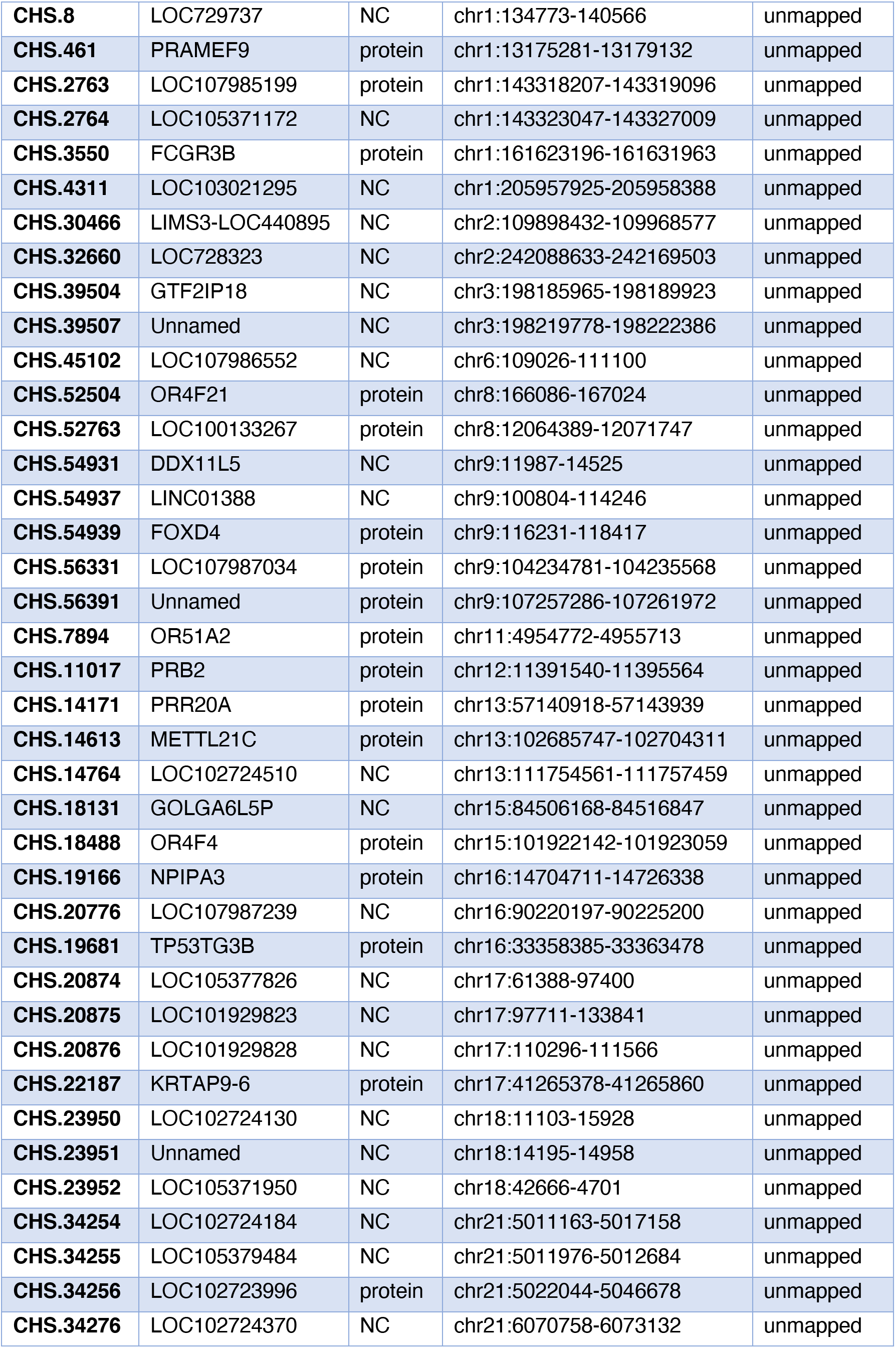
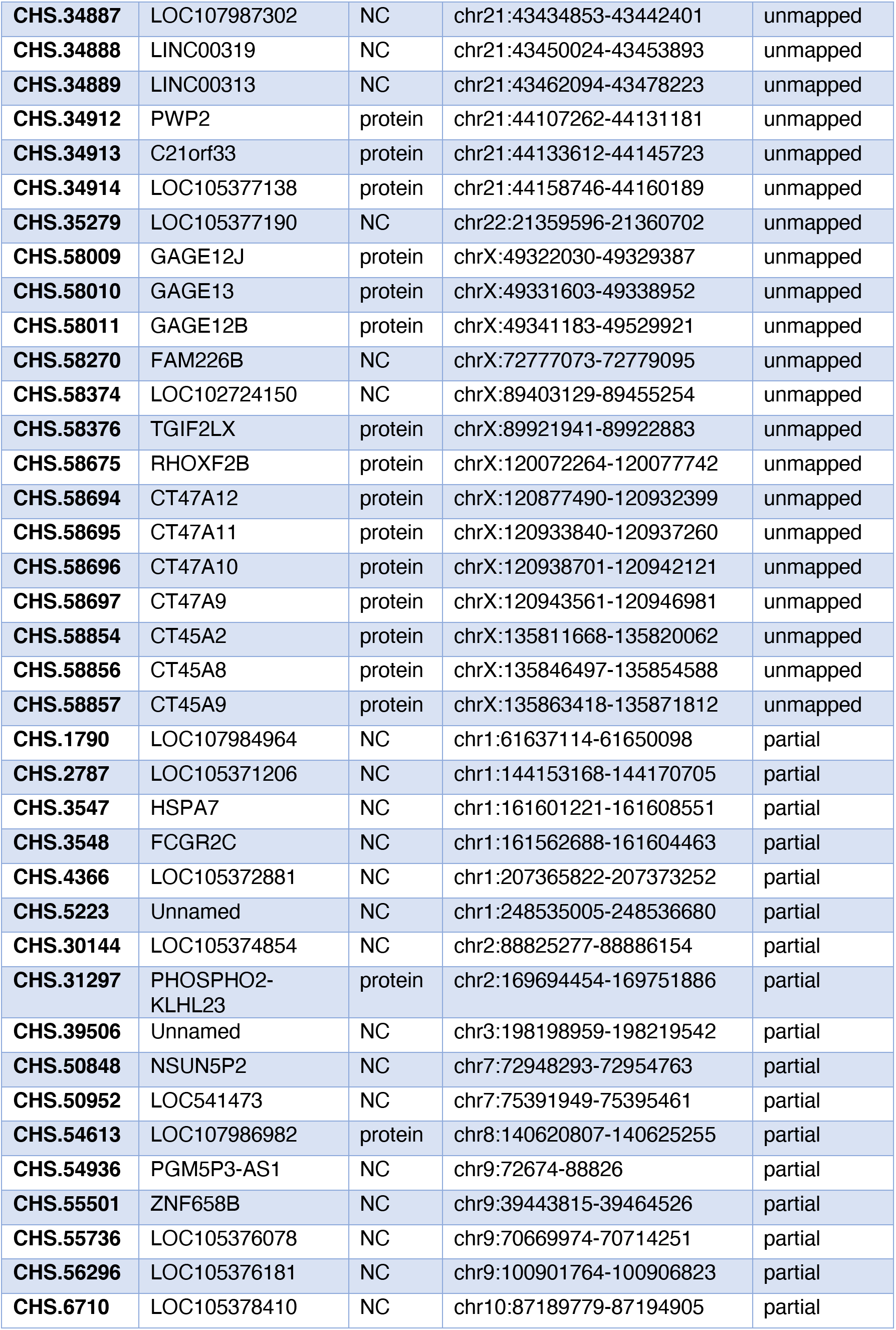
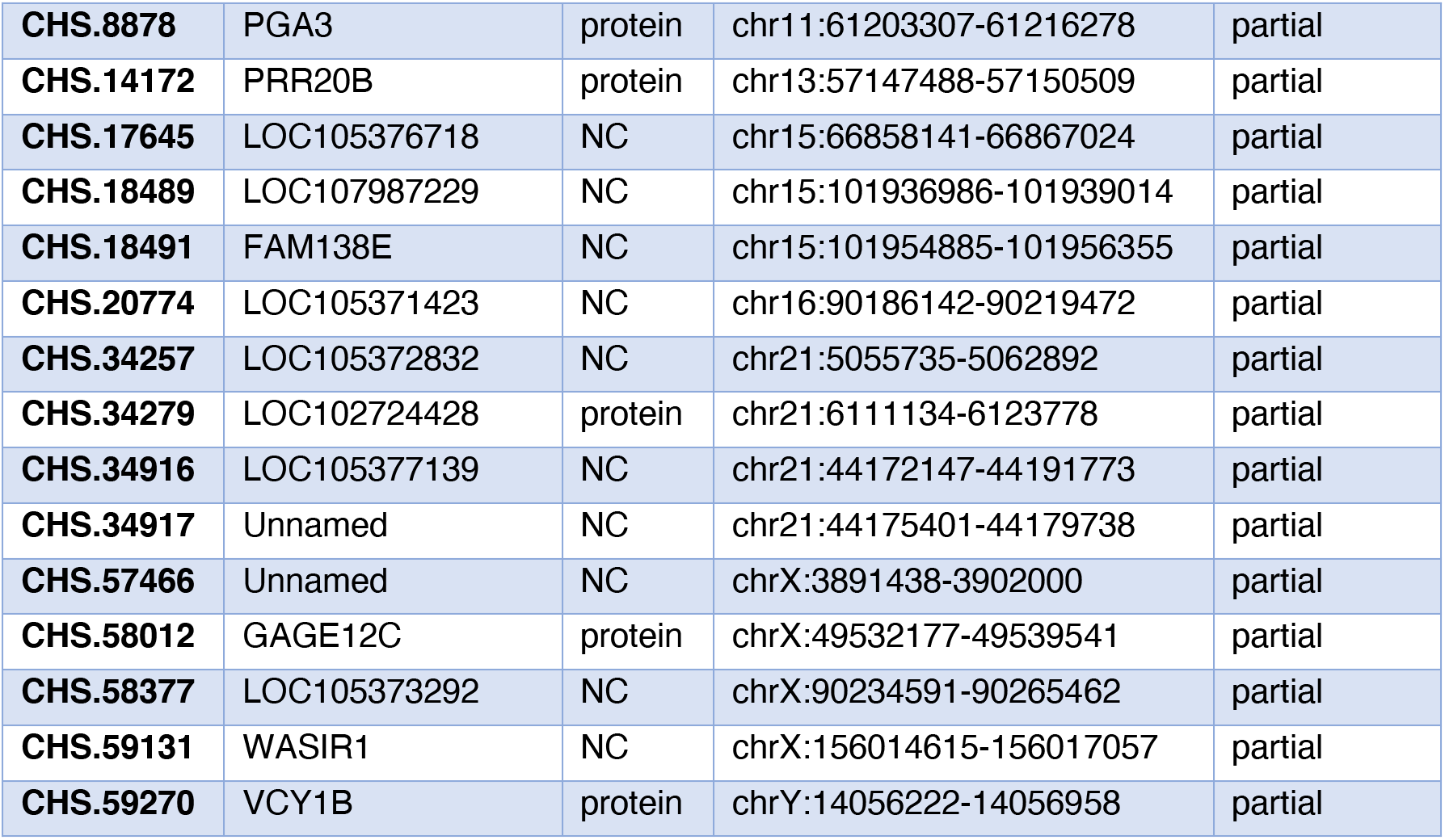
94 genes that are completely or mostly missing in Ash1. The Mapping status column shows “unmapped” if the gene is entirely missing from Ash1, and “partial” if less than 50% of the gene appears in Ash1. 40 of the genes are protein-coding and 54 are noncoding. All of the protein-coding genes are members of multi-gene families. Abbreviations: NC, noncoding.

After mapping the genes onto Ash1, we extracted the coding sequences from transcripts that mapped completely (coverage equal to 100%), aligned them to the coding sequences from GRCh38, and called variants relative to GRCh38 (see Methods). Within the 35,513,365 bp in these protein-coding transcripts, we found 20,864 single-nucleotide variants and indels. 14,499 of these variants fell within the GIAB “callable” regions for high-confidence variants, although 3,963 of these were in GIAB “difficult” repetitive regions, for which alignments are often ambiguous. Of the 10,536 variants not in these difficult regions, 10,456 (99.2%) agreed with the GIAB high-confidence variant set. In the difficult regions, 3,804/3,963 (96.0%) agreed with the GIAB set.

**Table 6.**
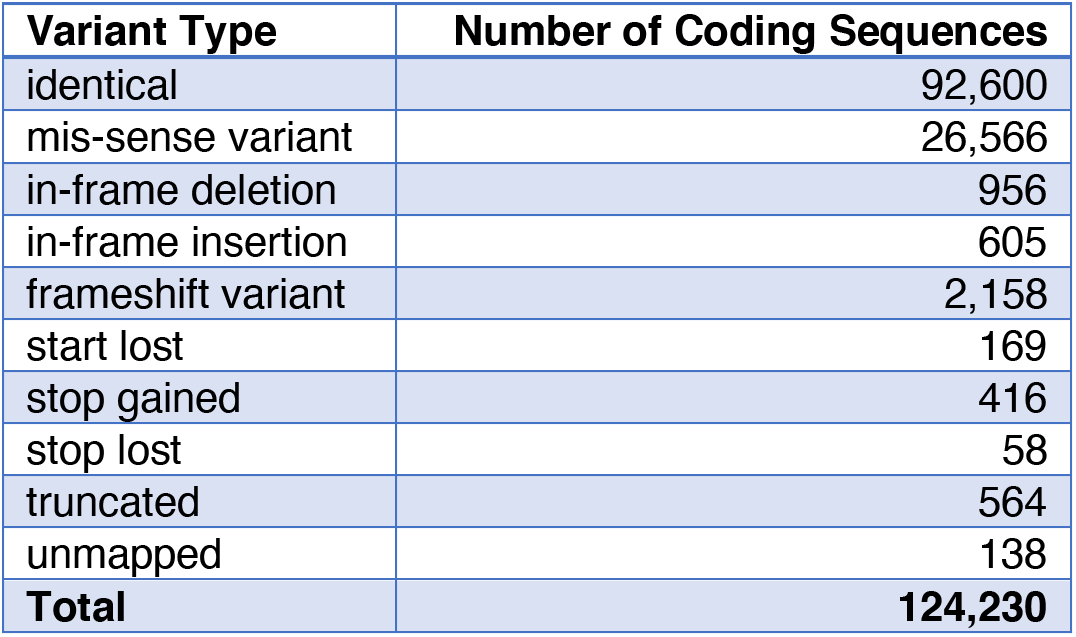
Comparison of protein coding sequences between Ash1 and GRCh38. Here, “insertion” means an insertion in Ash1 relative to GRCh38, and other terms are similarly referring to changes in Ash1 compared to GRCh38. “Truncated” indicates the transcript was only partially mapped. “Stop gained” refers to premature stop codons caused by a SNP.

| **Variant Type** | **Number of Coding Sequences** |
| --- | --- |
| identical | 92,600 |
| mis-sense variant | 26,566 |
| in-frame deletion | 956 |
| in-frame insertion | 605 |
| frameshift variant | 2,158 |
| start lost | 169 |
| stop gained | 416 |
| stop lost | 58 |
| truncated | 564 |
| unmapped | 138 |
| **Total** | **124,230** |

We then annotated the changes in amino acids caused by variants and incomplete mapping for all proteincoding sequences. Out of 124,238 protein coding transcripts from 20,197 genes, 92,600 (74.5%) have 100% identical protein sequences. Another 26,566 (21.4%) have at least one amino acid change but yield proteins with the identical length, and 1561 (1.3%) have frame-preserving mutations that insert or delete one or more amino acids, leaving the rest of the protein unchanged. **Table 6** shows statistics on all of the changes in protein sequences. If a protein had more than 1 variant, we counted it under the most consequential variant; i.e., if a protein had a missense variant and a premature stop codon, we include it in the “stop gained” group.

Of particular interest are those transcripts with variants that significantly disrupt the protein sequence and may result in loss of function. These include transcripts affected by a frameshift (2158), stop loss (58), stop gain (416), start loss (58), or truncation due to incomplete mapping (564). These disrupted isoforms represent 885 gene loci; however, 505 of these genes have at least 1 other isoform that is not affected by a disrupting variant. This leaves 380 genes in which all isoforms have at least one disruption; the full list is provided in Supplementary Table 1.

## Discussion

The assembly and annotation of this first Ashkenazi reference genome, Ash1, are comparable in completeness to the current human reference genome, GRCh38. We began by creating a high-quality de novo assembly of Ash1, using reads generated by multiple sequencing technologies, and then improved the assembly in multiple ways, using GRCh38 for chromosome-scale scaffolding and then using high-quality variant benchmarks from GIAB, computed on data from the same individual, to correct thousands of small consensus sequence errors. Unlike GRCh38, which represents a mosaic of multiple individuals, Ash1 is derived almost entirely from a single individual. More precisely, Ash1 v1.7 contains 2,973,118,650 bp mapped onto chromosomes, of which 98.04% derive from a single Ashkenazi individual, and the remaining 58,317,846 bp (1.96%) were taken from GRCh38. As more data and better assemblies become available, we expect this latter portion to shrink.

The gene content of Ash1 is nearly identical to GRCh38: all of the genes are present, with the only differences being 40 protein-coding genes and 54 noncoding genes (0.22% of the total) that are present in fewer copies. 11 genes were mapped to different chromosomes, suggesting a small number of chromosomal rearrangements that predominately involve exchanges of subtelomeric regions. It is likely that Ash1 contains additional copies of some genes, but we did not attempt to search for these.

Similarly to GRCh38, Ash1 is not yet complete, and we plan to improve the assembly over time, much as GRCh38 has improved since its initial release in 2001. Newer sequence data including ultralong reads (over 100,000 bp in length) have recently been generated, which should allow additional gap filling and polishing of the genome sequence. Although the estimated quality of Ash1 v1.7 is very high, some disagreements between the current assembly and the GIAB benchmarks remain, indicating further room for improvement, especially in the resolution of complex repetitive regions. Additional analysis may also be needed to confirm that the small number of missing and disrupted genes are genuine differences between the genomes rather than incorrectly assembled repeats.

Nonetheless, the Ash1 genome provides a ready-to-use reference for any genetic studies involving individuals with an Ashkenazi Jewish background. In these individuals, alignments to Ash1 should yield fewer variants than alignment against GRCh38, which in turn will allow investigators to spend less time eliminating irrelevant variants. In addition, the computational methods used in this study provide a recipe that should allow the construction of many more human reference genomes, representing the many different populations of humans in the world today.

## Methods

For the initial assembly of the combined Illumina, ONT, and PacBio data, we used MaSuRCA v3.3.4 (Zimin et al. 2013) to generate a set of contigs that we designated the Ash1 v0.5 assembly. We filtered the primary assembly for haplotype duplications by aligning the assembly to itself, and looking for contigs that were completely covered by other, larger contigs and that were >97% identical to the larger contig. This process filtered out 3,438 small contigs containing 56,956,142 bp. To assign the contigs to chromosomes, we used a scaffolding script included in MaSuRCA (chromosome_scaffolder.sh) that first aligned the assembly to the GRCh38.p12 reference genome using MUMmer4 (Marcais et al. 2018). Many contigs aligned end-to-end to a single chromosome, and these were easy to place. The script then considered the contigs that aligned to GRCh38 in multiple disjoint chunks. Each alignment that ended within a contig, and that was >5kb from either end of the contig, was designated a potential breakpoint.

The scaffolding script then aligned the ONT reads to the Ash1 v0.5 contigs using blasr (Chaisson and Tesler 2012) and computed the read coverage. A potential breakpoint was deemed a mis-assembly if there was a region of either low (<=3x) or high (>35x) coverage within 50 kb of the alignment breakpoint. This procedure identified 470 breakpoints and then split the Ash0.5 contigs at those mis-assemblies. Note that if a mis-assembly occurred in a low coverage region, the contig was split at the weak point. If the mis-assembly occurred in a high-coverage region, then it was likely due to a repetitive sequence, and the contig was split at the alignment breakpoint location. After splitting, the script re-aligned the split contigs to the GRCh38 reference and used the best alignments to assign each contig or partial contig to a chromosome location. The resulting “tiled” assembly, Ash1 v0.9, had 2,843,368,711 bases in 1,026 contigs assigned to specific chromosomes. The remaining contigs were left unplaced.

Some gaps in the initial Ash1 assembly occurred in areas where GRCh38 is ungapped, sometimes corresponding to regions that were manually curated to capture especially difficult repetitive regions. To capture these regions, we took two additional gap-filling steps. First, for every gap in Ash1 v0.9, we identified cases where contiguous GRCh38 sequence spanned the gap, with at least 2kb of GRCh38 aligning uniquely to Ash1 v0.9 on both sides of the gap. In these cases we filled the gap in Ash1 with the GRCh38 sequence. This step closed 412 gaps, yielding Ash1 v1.0. (Note that in the Ash1 genome, these GRCh38 sequences are recorded in lowercase, to distinguish them from the Ashkenazi-origin sequence, which is in uppercase.) Next, for the gaps where we could not find contiguous GRCh38 sequence that aligned to both sides of the gap in Ash1 v0.9, we looked for GRCh38 contigs that might fit into the gap, given the gap size estimate and the implied gap coordinates on GRCh38. We then inserted GRCh38 contigs that “fit” into the gaps surrounding them, leaving a 100bp gap (represented as 100 N’s) on both sides. This second step added 948 sequences from GRCh38 into the gaps, making the gaps smaller but leaving a pair of 100-bp gaps for each inserted contig. Some of these sequences were separate, small contigs in GRCh38, and some were derived from GRCh38 contigs that extended into gaps in Ash1 (see Suppl. Figure S1). This assembly, Ash1 v1.1, contained 948 more gaps than Ash1 v1.0, but the gaps were smaller. Overall, these two gap-filling steps added 58,317,846 bp of sequence from GRCh38.

We next searched Ash1 v1.1 for the 12,745 high-quality, isolated structural variants (insertions and deletions ≥50 bp) that Zook et al. identified by comparing the Ashkenazi trio data to GRCh37 (Zook et al. 2019). Because Ash1 has a different coordinate system from GRCh38, we had to use sequence alignment methods to find these SVs in Ash1. For this step, we extracted a region of sequence extending 1000 bp beyond each SV in both directions. (Note that if a variant occurred in a tandem duplication longer than 1000bp, this strategy might fail to align it to the correct location.) We then aligned each region to Ash1 using nucmer (Marcais et al. 2018), and filtered the results to determine which SVs were present and which were missing from Ash1 v1.1.

We also made small variant calls from Ash1 v1.1 relative to GRCh37, and compared these to the v4.0 benchmark variants from GIAB (which uses GRCh37) using the Global Alliance for Genomics and Health (GA4GH) Benchmark tools (Krusche et al. 2019). Our definition of a false positive variant (FP) included all variants from Ash1 not in the GIAB v4.0 set of variant calls (i.e., in the vcf file) but within the v4.0 regions, as well as variants from Ash1 not matching the v4.0 genotype; e.g., heterozygous variants in the benchmark that are homozygous variants from Ash1 because Ash1 represents only one haplotype. To ignore errors due to Ash1 representing a single haplotype and identify potential errors in Ash1, we excluded FPs where the v4.0 call was heterozygous or compound heterozygous (reported as FP.gt by the GA4GH benchmarking tools) or where the FP was within 30 bp of a v4.0 variant (reported as FP.al). To identify candidates for correction in the assembly, we also excluded FPs in UCSC GRCh37 vs. GRCh37 self-chain alignments longer than 10 kb, since these were potential collapses in the assembly that would need to be corrected in a different way. Using the remaining FPs, we corrected 32,814 substitution errors, 6670 insertion errors, and 14,151 deletion errors in the Ash1 assembly. This did not correct any regions in Ash1 that aligned outside the v4.0 benchmark regions for GRCh37. These corrections yielded Ash1 v1.2.

To create Ash1 v1.3, we added 2,786,257 bases to the beginning of the X chromosome and 2,281,641 bases to the beginning of the Y chromosome, based on careful alignments to GRCh38. These sequences, which are part of the pseudo-autosomal regions, are nearly identical between X and Y in GRCh38 and in Ash1. We also identified ~3 Mbp of sequence in two contigs that the assembler had labelled as “degenerate” that was missing from Ash1 but present on GRCh38, and we placed these contigs onto chromosomes.

To create v1.4, we realigned Ash1 v1.3 to GRCh38 using more sensitive parameters, allowing us to place a few additional contigs onto chromosomes. We then re-polished the v1.4 assembly with the POLCA software (Zimin and Salzberg 2019) to reduce the number of errors in the consensus, applying polishing to all of the sequences added in previous refinement steps. These steps made 54,125 substitution corrections and corrected 264,165 bases in insertion/deletion errors, yielding Ash1 v1.6.

Finally, in our initial comparison to the gnomAD Askhenazi major alleles, we found 273,866 heterozygous SNV sites where the GRCh38 reference allele appeared in the Ash1.6 assembly but where HG002 contained the Ashkenazi major allele as well. For these sites, we replaced the Ash1 reference allele with the Ashkenazi major allele. Because the initial assembly arbitrarily chose a representative base at heterozygous sites, this step preserved the assembly’s fidelity to the underlying HG002 sequence. These single-base changes resulted in Ash1 v1.7.

### Unplaced contigs

After chromosome assignment was done, 947 contigs remained unplaced. From those, we identified 92 contigs containing 5,118,131 bp as centromeric repeats; 26 contigs containing 5,716,977 bp mapped to unplaced scaffolds in GRCh38.p12, and the remaining 829 contigs containing 42,641,604 represent additional unknown contigs. All 829 unplaced contigs have their best matches to other human sequence, with alignment identities ranging from 78–97%.

### Aligning long PacBio reads for validation

We downloaded a recently released set of PacBio HiFi reads, generated on the Sequel II System, from the HG002 Data Freeze (v1.0) at Human Pangenome Reference Consortium github site (https://github.com/human-pangenomics/HG002_Data_Freeze_v1.0#hg002-data-freeze-v10-recommended-downsampled-data-mix). These data, which were not used in our assembly of Ash1, were size selected for 15 Kb and 20 Kb fragments, and they represent ~34x genome coverage of the genome. We aligned them to Ash1 v1.7 genome using bwa-mem with default parameters. We filtered the alignments using samtools to include only reads aligning with quality of 40 and above.

### Benchmarking Ash V1.6 Against GIAB HG002 V4.1 Benchmark set

Variant calls for Ash V1.6 assembly against the GRCh38 reference without alternate loci or decoy sequences (available from ftp://ftp.ncbi.nlm.nih.gov/genomes/all/GCA/000/001/405/GCA_000001405.15_GRCh38/_seqs_for_alignment_pipelines.ucsc_ids/GCA_000001405.15_GRCh38 no alt analysis set.fna.gz) were made using dipcall version 0.1 (Li et al. 2018). The resulting variant calls were compared to GIAB HG002 V4.1 benchmark set using the hap.py benchmarking tool (Krusche et al. 2019).

Because the assembly represents a single haplotype, FPs were calculated differently from the standard hap.py output, where FPs due to genotype and allele mismatches were subtracted from the total false positives. QV values were calculated using the modified FP metric, QV = −10*log(FPs/benchmark region size), where benchmark region size was “Subset.IS_CONF.Size” from the hap.py output.

### Mapping gnomAD SNVs onto Ash1

For each of the 2,008,397 gnomAD SNV sites where the Ashkenazi major allele differed from GRCh38, we extracted a 2kb region centered on the SNV from GRCh38. We aligned these 2kb sequences using nucmer (Marcais et al. 2018) with a requirement that seed matches be at least 50 bases (-l 50) and that anchors be unique in the reference and query (--mum), to help eliminate spurious mappings in repetitive regions, though this reduced the number of SNVs considered. We then filtered the alignments further using delta-filter to collect alignments spanning at least 1980 bases (-l 1980) with at least 99% identity (-i 99), and took the best alignment of each region (-q). Coordinates were then converted to Ash1 by using the delta2paf utility from paftools (Li 2018), followed by paftools liftover on the paf file to obtain the Ash1 genome coordinates of each original SNV site. This process unambiguously mapped 1,790,688 SNVs (89%) onto Ash1.

### Comparing variants in mapped genes

Gffread was used to extract the coding sequences from GRCh38 and Ash1. Sequences were aligned pairwise using the EMBOSS Stretcher command line interface (Madeira et al. 2019) from Biopython 1.75. The alignments were used to determine the GRCh38 location, sequence, and functional consequence of each variant. When comparing GIAB HG002 V3.3.2 benchmark set, we excluded any transcripts that did not map with an alignment coverage of 100%. We compared the variants to the benchmark set using vcfeval from RealTimeGenomics tools (Cleary et al. 2015). We used bedtools (Quinlan and Hall 2010) to intersect the false positive variants in Ash1 genes with the union set of difficult regions from GIAB (ftp://ftp-trace.ncbi.nlm.nih.gov/giab/ftp/release/genome-stratifications/v2.0/GRCh38/union/GRCh38_alldifficultregions.bed).

### Aligning transcripts between GRCh38 and Ash1

To compute the cumulative distributions shown in Supplemental Figs. 2 and 3, the mRNA sequences of the Ash1 transcripts and GRCh38 transcripts were extracted using gffread. The sequences were then aligned pairwise using the EMBOSS Stretcher command line interface (Madeira et al. 2019) from Biopython 1.75, and the resulting alignments were used to calculate the percent of GRCh38 transcript lengths covered and the percent identity between the pairs of transcripts.

### Data availability

The Ash1 assembly, including current and earlier versions, is freely available at https://github.com/AshkenaziGenome/Assembly and has been deposited in GenBank as accession GCA_011064465.1 and BioProject PRJNA607914. The github site also contains the gene annotation and an index with a mapping between the identifiers used by CHESS, RefSeq, and GENCODE.

## Acknowledgements

Thanks to Adam Phillippy for helpful comments on drafts of this manuscript. This work was supported in part by NIH under grants R01-HG006677 and R35-GM130151, and by the USDA National Institute of Food and Agriculture under grant 2018-67015-28199. Certain commercial equipment, instruments, or materials are identified to specify adequately experimental conditions or reported results. Such identification does not imply recommendation or endorsement by the National Institute of Standards, nor does it imply that the equipment, instruments, or materials identified are necessarily the best available for the purpose.

## Supplementary Figures

**Figure S1.**
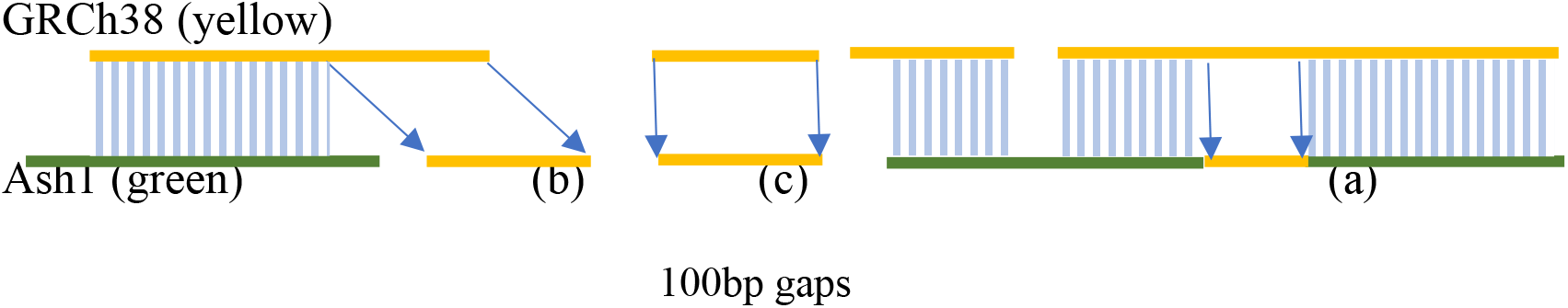
Adding sequences from GRCh38 to the Ash1 genome assembly. In (a), GRCh38 closes a gap in Ash1. In (b), the GRCh38 contig extends into a gap in Ash1, but the sequence adjacent to the gap does not match. If the GRCh38 extended >1000bp into the gap, and if the alignment ended > 100bp from the end of the Ash1 contig, then the GRCh38 sequence indicated by (b) was inserted, separated from the Ash1 sequence by a gap set to 100 Ns. Case (c) shows an example where a separate GRCh38 contig falls completely within a gap in Ash1, in which case it would be inserted with gaps on both sides.

**Figure S2:**
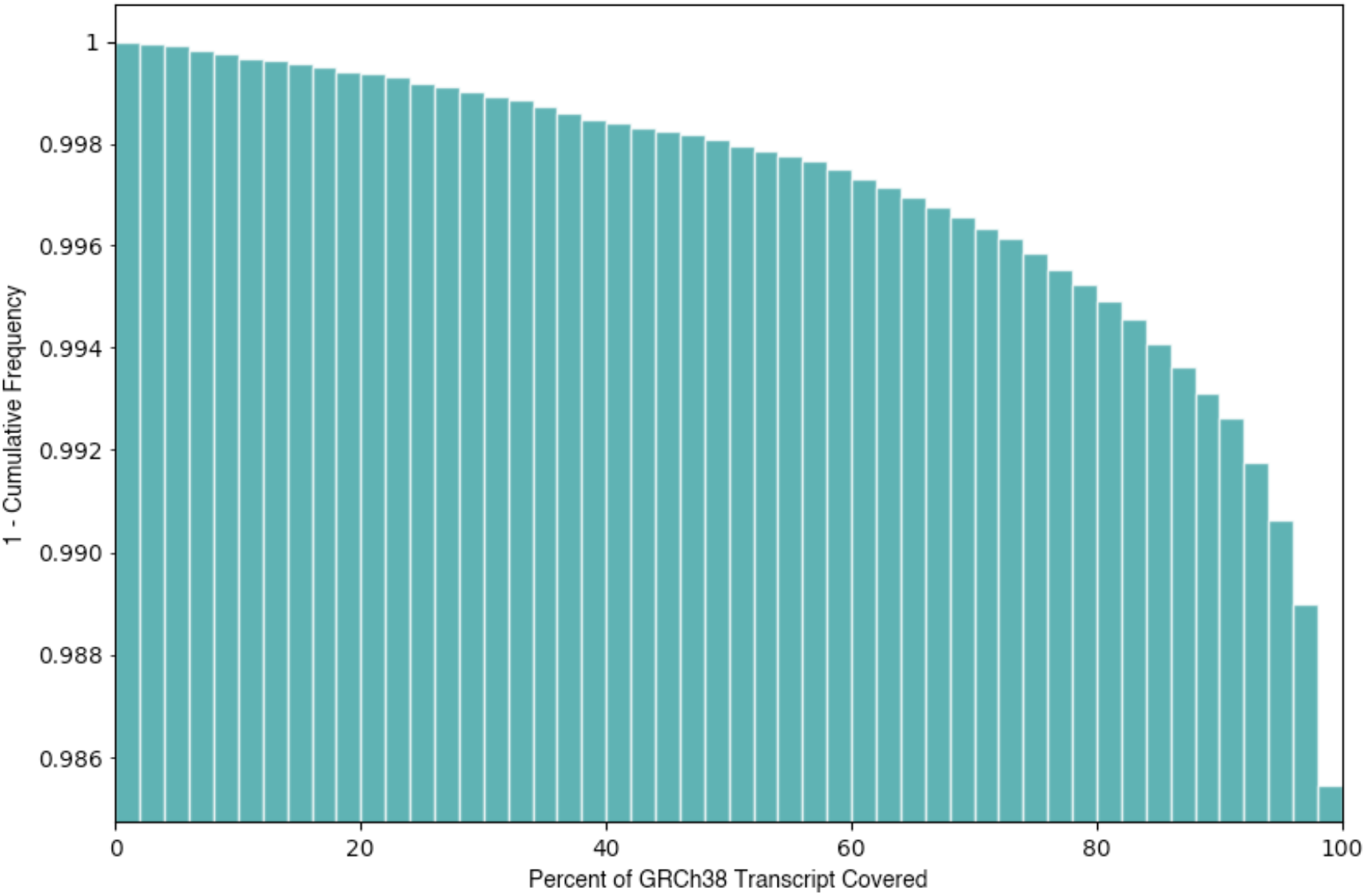
Cumulative distribution showing how much of the GRCh38 transcripts map onto Ash1. The Y axis shows the fraction of transcripts with percent coverage greater than or equal to coverage on the X axis; e.g., the next-to-last bar at 98% on the X axis shows that 98.9% of GRCh38 transcripts (Y axis) mapped for at least 98% of their length onto Ash1.

**Figure S3:**
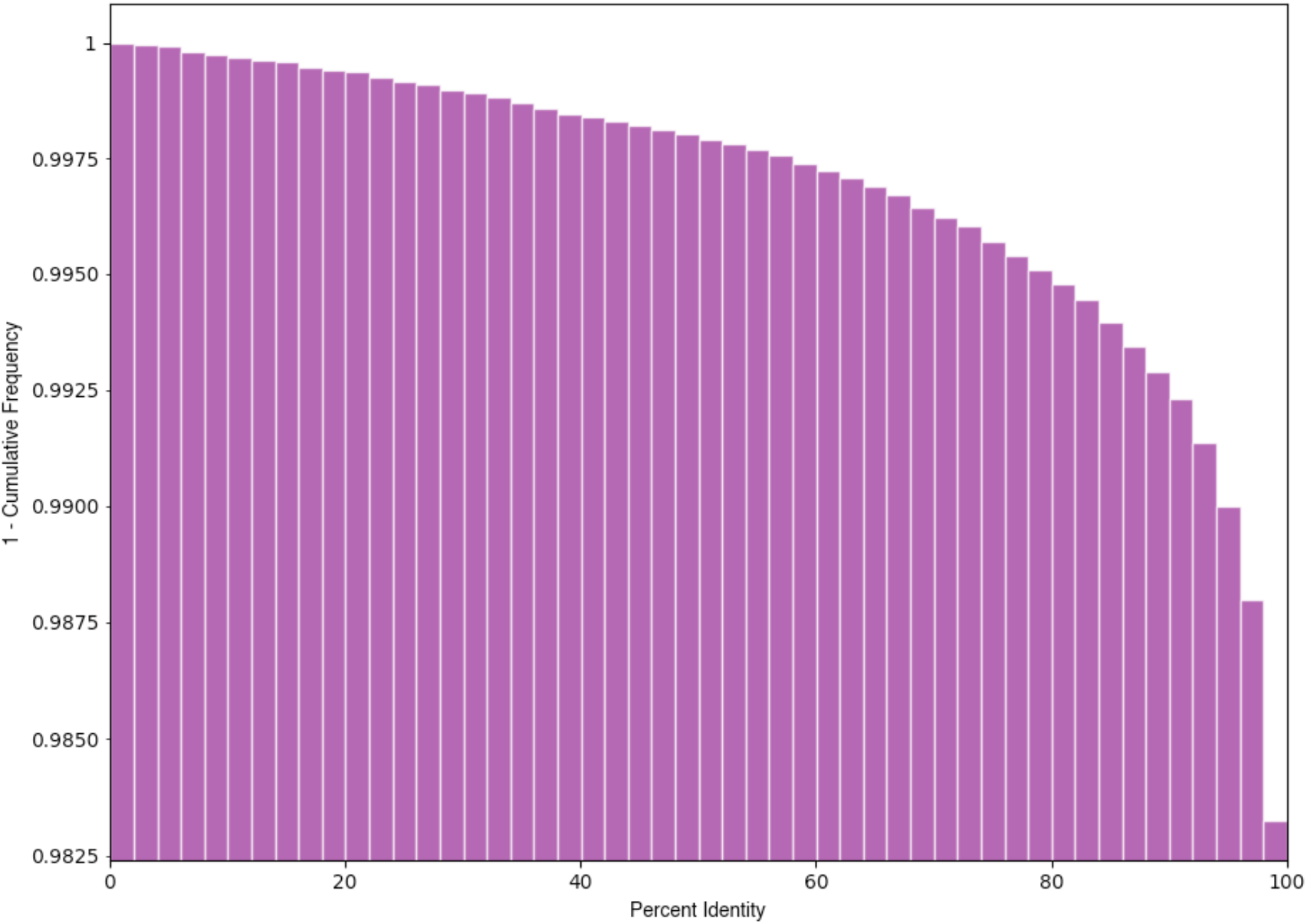
Cumulative distribution of the sequence identity of transcripts mapped onto Ash1. The Y axis shows the fraction of transcripts that aligned between GRCh38 and Ash1 with DNA sequence identity greater than or equal to the percent identity on the X axis. E.g., the next-to-last vertical bar at 98% on the X axis shows that 98.75% of the GRCh38 transcripts aligned at 98% or greater identity to Ash1.

## Notes

https://github.com/AshkenaziGenome

